# nsearch: An open source C++ library for processing and similarity searching of next-generation sequencing data

**DOI:** 10.1101/399782

**Authors:** Schmid Steven, Julian Timothy R., Tamminen Manu

**Affiliations:** Department of Biosystems Science and Engineering, ETH Zürich, Zürich, Switzerland; Eawag, Swiss Federal Institute of Aquatic Science and Technology, Duebendorf, Switzerland; Swiss Tropical and Public Health Institute, Basel, Switzerland; University of Basel, Basel, Switzerland; Department of Biology, University of Turku, Turku, Finland

**Keywords:** nsearch, C++11, paired-end read merging, quality filtering, sequence similarity searching

## Abstract

Advancements in DNA sequencing technologies rapidly change the landscape of modern biology. The novel next-generation sequencing (NGS) applications often have special requirements regarding experimental data processing. Software tools developed and used for novel applications are generally designed for specific use cases and as such may be difficult to adapt to new uses. Simultaneously, software tools designed to be general are often difficult to adapt to special use cases.

Here, we present nsearch, a modern open source C++11 library and command-line tool for biological sequence data processing. nsearch offers commonly used components for handling of biological sequences including paired-end read merging, quality filtering and sequence similarity searching. nsearch can either be embedded natively into other C++ applications or be packaged as a standalone executable.

Functionality and performance of nsearch is shown using benchmark data created using the Rfam 13 database. Benchmarking against common general purpose tools USEARCH and VSEARCH demonstrates that nsearch delivers performance comparable these state-of-the-art tools.

nsearch is available on GitHub under the permissive BSD-3-clause license: https://github.com/stevschmid/nsearch

## Introduction

DNA sequence data is being generated at an increasing speed due to ongoing innovation in next-generation sequencing (NGS) technologies. This trend requires bioinformatic tools that are capable of handling the increasing sequence data volumes. Consequently, the landscape of next-generation DNA sequencing applications and related bioinformatic tools is expanding rapidly, resulting in a large variety of experimental approaches ranging from microbiome analyses (Caporaso et al., 2010) to transcriptional profiling (Dobin et al., 2013). Despite the diversity of these applications, certain basic operations such as raw sequence data processing and nucleotide database searches are universal. For example, BLAST-like nucleotide database searches are common features in bioinformatic applications. Nevertheless, easily customizable, embeddable implementations of such algorithms are currently not available. For instance, while USEARCH (Edgar, 2010) provides a fast, high-quality tool for nucleotide database searching, it is proprietary software and requires a license for processing large data volumes. The open-source alternative VSEARCH (Rognes et al., 2016) suffers from limited flexibility and reusability as a library.

Here, we present here nsearch, a high-performance, lightweight, customizable and embeddable NGS data processing toolkit. nsearch is a C++11 NGS data processing library of algorithms frequently encountered when handling, analyzing, or interpreting NGS data with no external dependencies. Its modular architecture offers functionality in form of embeddable components. The option to natively integrate nsearch eliminates the need to package it or list it as a prerequisite for end-user compatible bioinformatics software. nsearch can also be used as a standalone command line tool in the vein of USEARCH (Edgar, 2010) or VSEARCH (Rognes et al., 2016).

nsearch offers:

- Basic handling for biological sequences (access residues, create complement, reverse a sequence)
- File support for FASTA/FASTQ (compressed or uncompressed) and ALNOUT
- Merging of paired-end reads
- Quality filtering of reads
- Database searching (e.g. to map reads to a list of reference sequences)
- Support for DNA, RNA and amino acid sequences (full IUPAC codeset, Cornish-Bowden (1985))
- Unit tests for all components
- Permissive license without copyleft restriction: The BSD-3-clause license was chosen to maximize utility and encourage project participation

## Implementation

nsearch handles and processes data generated from short read sequencers. Functionality includes paired-end read merging, quality filtering, and database searching. Current implementation allows handling of both DNA (inclusive of DNA and RNA) and Protein alphabets, with policies implemented via C++ templates.

### Alphabets

To distinguish between different types of biological sequences, nsearch uses the concept of alphabets. Alphabets define how biological sequences and its letters are treated. nsearch currently supports two alphabets: DNA (nucleic acids, DNA and RNA) and Protein (amino acids).

The alphabets differ by their policies, which are implemented via C++ templates. The following policies describe an alphabet:

- BitMapPolicy: How the alphabet can be represented compactly as bits (e.g. DNA can be represented with 2 bits).
- ComplementPolicy: How letters of an alphabet can be complemented (e.g. A↔T for DNA).
- ScorePolicy: How two letters are scored to each other (default for DNA is +2/-4 for a match/mismatch, default for Protein is the BLOSUM62 (Henikoff and Henikoff, 1992) scoring matrix).
- MatchPolicy: If two letters can be considered identical (Boolean).

The C++ policy templates enable the straightforward implementation of custom alphabets or different scoring schemes. Nearly all components in nsearch require the developer to specify an alphabet. For example, FASTA::Reader<DNA> reads a FASTA file and returns DNA/RNA sequences as Sequence<DNA>. The explicit alphabet specification gives developers the option to change the default alphabet-specific behavior of algorithms at compile-time.

### Paired-End Read Merging

nsearch uses an exhaustive search strategy for finding the best overlap between two paired reads. This exhaustive search strategy has the advantage that it requires no prior information about the alignment of the paired reads. Evaluating every possible alignment is suitable for small reads in the hundred nucleotides range.

An alignment is checked for: a) that the length of the overlapping region is equal or above a fixed threshold (16nt), and b) that the identity in the overlap region is above a minimum identity threshold (90%). If no alignment fulfills these conditions, the forward and reverse reads are discarded.

If these conditions are met, the posterior error probabilities for this alignment are determined and used to score the alignment. The best scoring alignment will be used for the creation of the consensus read.

The consensus read is created by using the base calls of the forward and reverse read and by calculating the posterior quality scores in the overlapping region. In the overlapping region, the base call with the higher prior quality score is used. Agreeing base calls in the overlap increase the quality score, whereas disagreement leads to a lower quality score. Figure 1 illustrates the consensus building process. nsearch uses the formulas in Edgar and Flyvbjerg (2015) for calculating the posterior quality scores.

**Figure 1.**
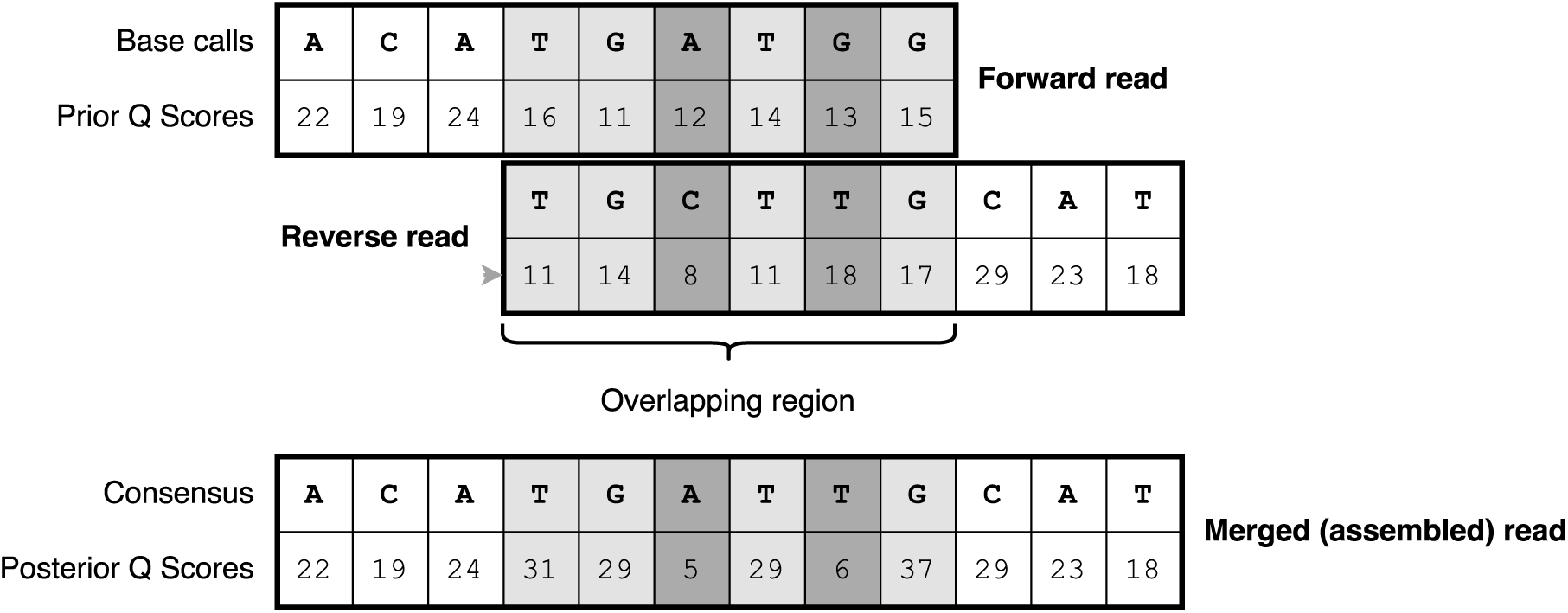
Determining the consensus sequence and posterior quality scores. Adapted from Edgar and Flyvbjerg (2015).

### Quality Filtering

The quality of a read is determined by the ability of a sequencer to determine the individual bases of a sequence unambiguously. A natural way to quantify the read quality is by the number of expected errors *E*. E is the sum of the error probability for each base call *p*_*i*_ (Edgar and Flyvbjerg (2015)), or:

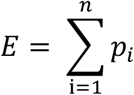

The error probabilities for each base call are based on the quality score as determined by the sequencing platform. The quality filtering of the nsearch command line tool discards reads for which the number of expected errors *E* exceeds a user-defined threshold (*E*_*max*_, default 1.0).

### Database Searching

nsearch also incorporates database searching based on the search strategy implemented in USEARCH (Edgar, 2010). Finding similar sequences in a reference database for a given input sequence (query) is a prominent and recurring problem for the analysis of biological sequences, including the mapping of reads. Due to the growing demand for faster algorithms to tackle the increase in sequencing throughput, new search tools such as USEARCH (Edgar, 2010) or VSEARCH (Rognes et al., 2016) emerged and promised “Search […] orders of magnitude faster than BLAST”.

High-throughput sequencing data requires a shift of the balance between sensitivity (i.e. finding all similar sequences) and run time (i.e. finding hits as fast as possible) towards the latter. The search strategy of USEARCH (Edgar, 2010) is based on the idea that a few good hits are sufficient for most sequence analysis problems. This restriction makes it possible to check only a limited number of candidate sequences. Candidate sequences are selected from the database based on the number of unique words (subsequences of length *k*, or k-mers) *U* they share. Candidates are checked in decreasing order of *U*: The sequence alignment and subsequently the sequence similarity is determined. If the similarity exceeds a user-defined threshold, the candidate is considered a hit. If the similarity is below the threshold, the candidate is considered a reject and the next best candidate is checked, until the maximum number of allowed rejects (trials) for this query has been reached. This early termination strategy makes sense, since the similarity decreases with each subsequent candidate (the number of unique shared words *U* also decreases, see Edgar (2004)). This heuristic, implemented here in nsearch, strikes a good balance between sensitivity and run time. The heuristic is useful for NGS data applications in that it allows fast searching while still maintaining sufficient sensitivity.

### Indexing

Database searching in nsearch starts with indexing. During the indexing step, the database sequences are stored in lookup tables. Lookup tables allow fast memory access to values for a given key. The search algorithm will use the lookup tables to quickly identify shared k-mers (Fig. 2). All k-mers of of every database sequence are stored for fast lookup. By default, nsearch uses *k* = 8 for the DNA alphabet and *k* = 5 for the Protein alphabet. Higher *k* leads in general to faster search run time at the price of reduced sensitivity. Within nsearch, each k-mer is transformed into a compact binary format (32-bit unsigned integer). The transformation is described by the BitMapPolicy of the alphabet (e.g. 2 bits are used to encode all DNA nucleotides).

**Figure 2.**
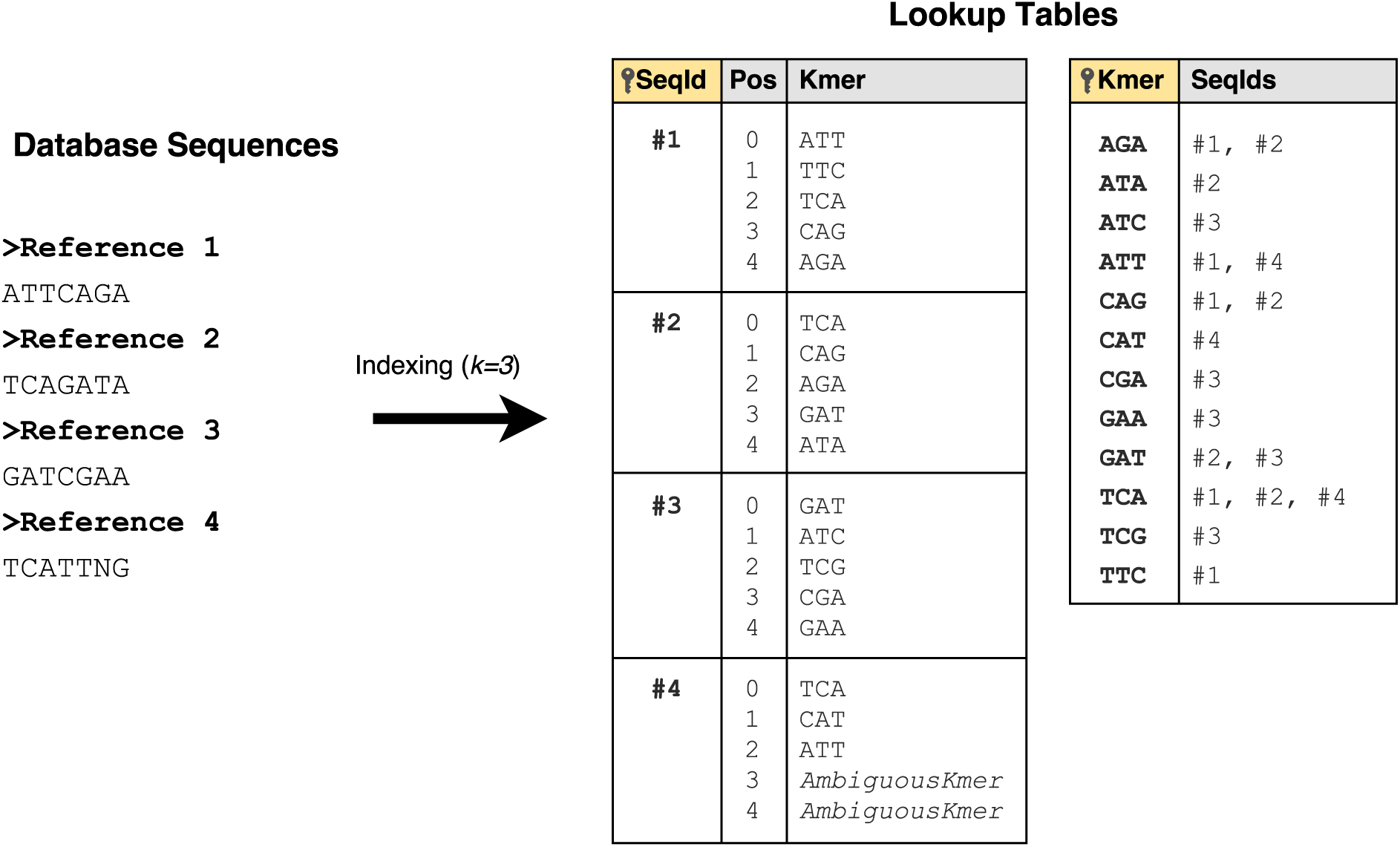
Database indexing. Indexing example for k-mers of length *k* = 3. During indexing, two lookup tables are built using the k-mers of every database sequence. One table (left) allows fast access to all k-mers of a given sequence. K-mers containing ambiguous residues (e.g. N for DNA) are treated as a special unique k-mer AmbiguousKmer. The second table (right) grants fast access to sequences which contain a given k-mer and is used in the candidate selection process.

K-mers which contain at least one ambiguous residue are mapped into a special k-mer called AmbiguousKmer. Storing each possible realization of an ambiguous k-mer is infeasible due to exponential growth. For example, a single DNA k-mer containing 5 N has 1024 different realizations. Storing all realizations of each k-mer would put a heavy burden on the memory consumption and would decrease the lookup efficiency. There is a trade-off between run time (faster lookup, less memory consumption) and sensitivity (being able to identify candidates which contain ambiguous residues). Due to the rarity of ambiguous nucleotides in NGS applications, regions containing ambigous nucleotides were discarded during database indexing.

### Searching

The search strategy was adopted from USEARCH (Edgar, 2010) and consists of two main steps: 1) Finding a list of promising candidates (based on the number of unique k-mers they have in common,*U*) and then 2) searching for hits among the high-scoring candidates by checking the sequence identity between query and candidate. A flowchart of the search strategy for a single query is shown in Fig. 3.

**Figure 3.**
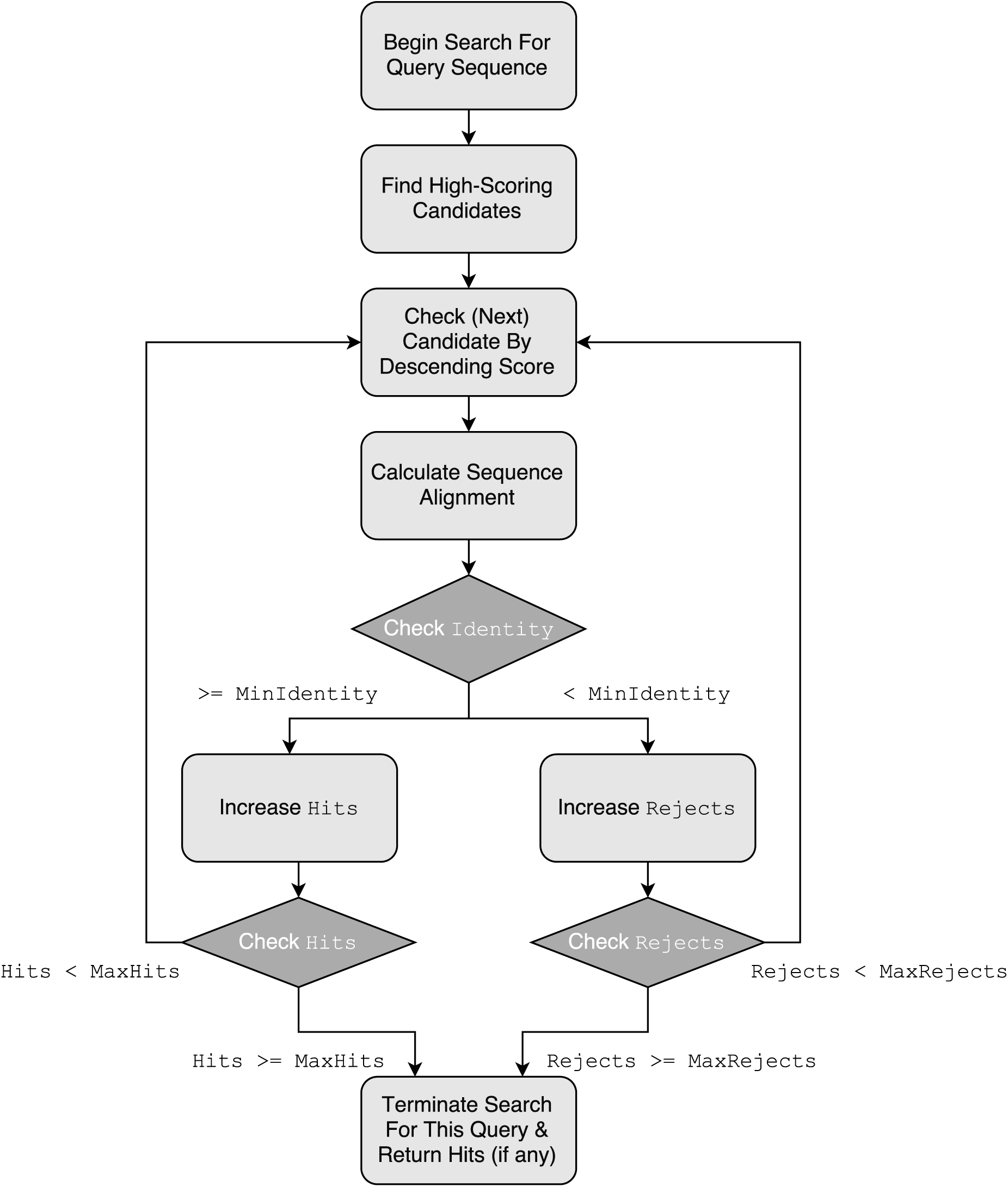
Flowchart visualization of the search strategy in nsearch for a single query. The search strategy was adopted from USEARCH (Edgar, 2010). The “Find High-Scoring Candidates”-step is outlined in Fig. 4. The “Calculate Sequence Alignment”-step is visualized in Fig. 5. Search for a query is terminated if either the maximum user-requested hits (default 1) or rejects (default 16) have been reached.

Based on the principle that similar sequences tend to have more k-mers in common (Edgar, 2004), the search algorithm uses the indexed database to search for candidate sequences by counting the number of unique k-mers they share with the query sequence (*U*). The candidate selection and high score building process is shown in Fig. 4.

**Figure 4.**
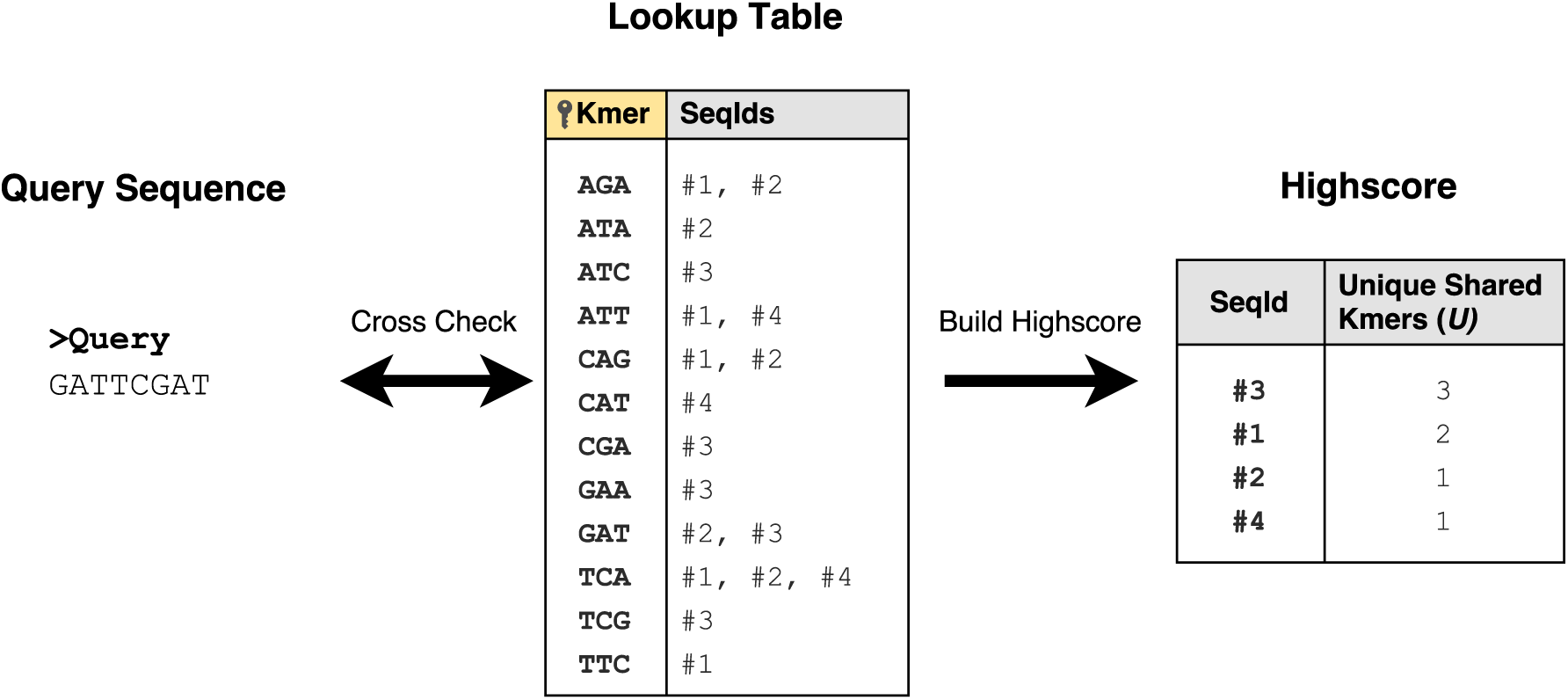
Candidate selection for a single query. Every k-mer (here *k* = 3) of the query sequence is checked against a lookup table built during database indexing. Each occurrence of a yet unencountered k-mer shared by both the query and a database sequence is counted (*U*). The unique counts *U* are used to build a sorted high score of candidates.

The sorted list of high-scoring candidates is then checked in decreasing order of *U*: The database sequence which shares the highest number of unique k-mers with the query sequence is checked first. For this purpose, the pairwise sequence alignment is determined. The sequence alignment is used to calculate the sequence identity, which in turn is used to determine whether the candidate is a hit.

First, the sequence alignment algorithm determines HSPs (high-scoring segment pairs) by employing a seed-and-extend strategy (Altschul et al., 1990). The HSPs are subsequently joined into a chain in a greedy manner: The highest scoring HSP is combined with the next-best non-overlapping HSP, until there are no HSPs left to join. The missing alignments between the HSPs are determined with banded DP (Chao et al., 1992). The sequence alignment process is visualized in Fig. 5.

**Figure 5.**
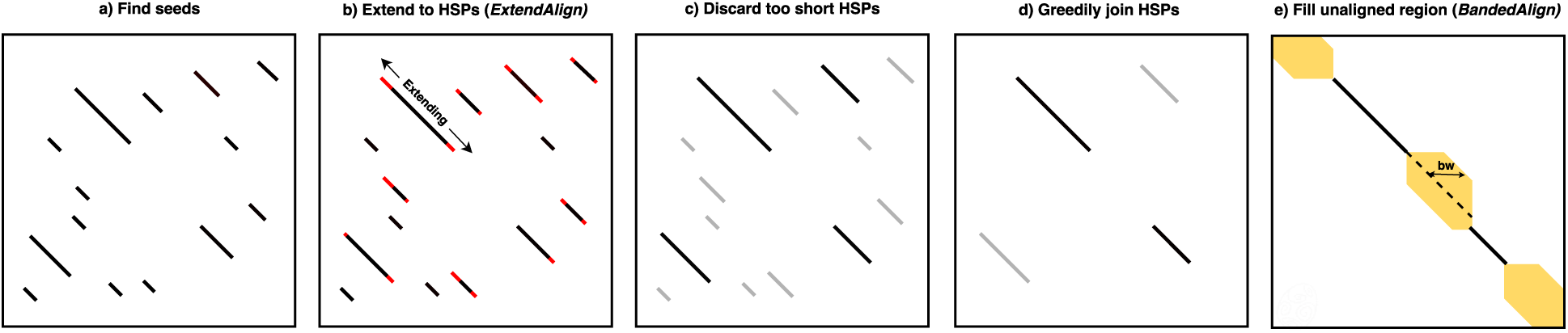
Sequence alignment. Shown is matrix visualization of the efficient global alignment process of nsearch in a matrix. a) The longest possible exactly matching subsequences (seeds) are determined. b) The seeds are extended using ExtendAlign, a BLAST -inspired gapped extension algorithm (Altschul et al., 1990), resulting in high-scoring segment pairs (HSPs). c) Short HSPs are discarded. d) Non-overlapping HSPs are greedily joined together. e) The unaligned regions between the HSPs are aligned using BandedAlign (banded DP, Chao et al. (1992)).

## Benchmarks

To assess the performance of nsearch, we employed a benchmark inspired by the original USEARCH publication (Edgar, 2010) and compared the database searching performance of the nsearch command line tool to USEARCH (Edgar, 2010) and VSEARCH (Rognes et al., 2016). These tools were selected because they are based on the same search strategy (termination after a maximum number of rejects) and thus can be compared directly.

The benchmark data set was created using the Rfam 13.0 database (Kalvari et al., 2018). The Rfam database is an annotated collection of non-coding RNA (ncRNA) families and is based on multiple sequence alignments. For every family which contains 2 or more sequences, one sequence was selected randomly and put in the query set. The remaining sequences of the family were added to the reference database, resulting in a query set of 2,686 sequences and a reference database of 2,312,296 sequences.

The following search settings were used: (1) report one hit (which has at least a 75% sequence identity between query and database sequence), (2) terminate after 8 rejects (high scoring candidates which have below 75% sequence identity), (3) use all available CPU cores, (4) no masking and (5) hardware: macOS High Sierra 10.13.4, Intel Core i7-3820, 16 GB Memory.

The benchmark objective was to find a member originating from the same family as the query sequence. The results are listed in Table 1.

**Table 1.**
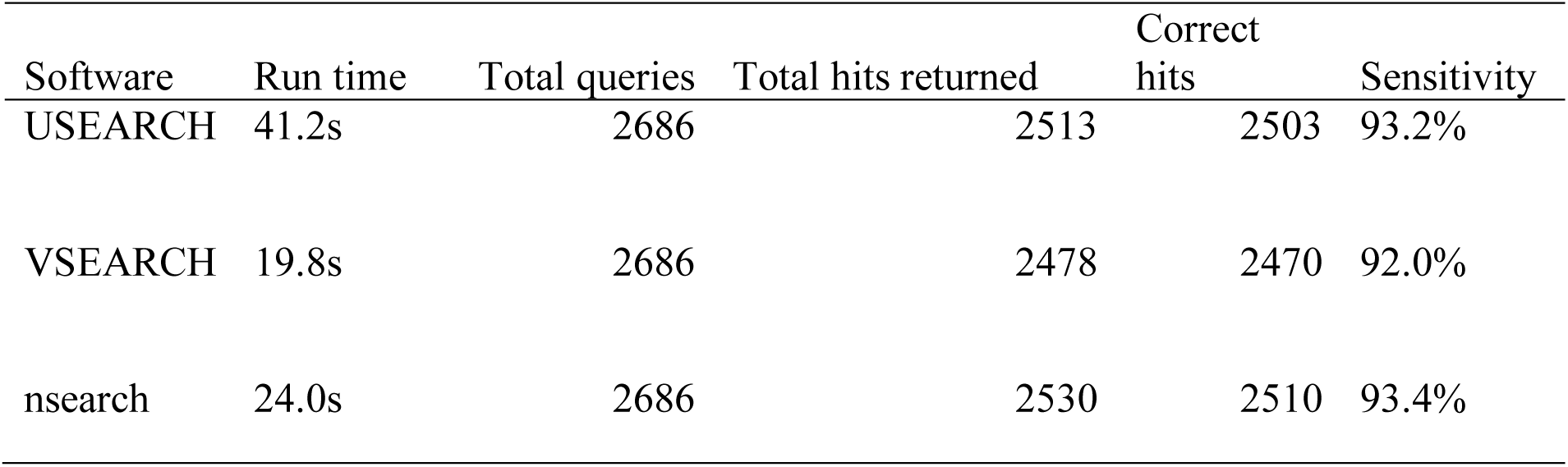
Search performance comparison of nsearch to the popular sequence analysis tools USEARCH and VSEARCH. The Rfam 13.0 database was used to create a benchmark data set with 2,686 query sequences and 2,312,296 database sequences. Each query sequence was extracted from a unique family of sequences. Correct hits are hits where the query and retrieved database sequence share the same family. The sensitivity is calculated as the number of correct hits divided by the number of total queries.

| Software | Run time | Total queries | Total hits returned | Correct hits | Sensitivity |
| --- | --- | --- | --- | --- | --- |
| USEARCH | 41.2s | 2686 | 2513 | 2503 | 93.2% |
| VSEARCH | 19.8s | 2686 | 2478 | 2470 | 92.0% |
| *nsearch* | 24.0s | 2686 | 2530 | 2510 | 93.4% |

Since all 3 tools share the same search strategy, we suspect that the difference in run time performance can be mainly attributed to the sequence alignment approach. VSEARCH (Rognes et al., 2016) calculates the full DP matrix with SIMD vectorization and can capitalize on the short sequences used in this benchmark. Due to USEARCH (Edgar, 2010) being a proprietary software (closed source), it’s difficult to explain the stark performance difference between USEARCH (Edgar, 2010) and nsearch. Both are using heuristics for the sequence alignment process. nsearch is using a seed-and-extend strategy (Altschul et al., 1990) in combination with banded DP (Chao et al., 1992).

*nsearch* performance was directly comparable with USEARCH and VSEARCH, with a run time approaching VSEARCH, and the highest sensivity (Table 1).

## Summary

nsearch is a zero-dependency C++ software library that offers components and algorithm for short read DNA sequencer data processing. It provides modular components for biological sequence handling, currently implemented to handle both DNA and protein alphabets, paired-end read merging, quality filtering and database searching. nsearch can be embedded natively into other bioinformatics software. The choice of developing nsearch as a library makes wrappers for other programming languages feasible. This would allow the incorporation in other programming language environments (such as Python or Ruby).

Developers can combine and fine-tune components of nsearch to tackle novel data processing use cases. nsearch reduces the time for researchers to add native NGS data processing capabilities to their own software. nsearch can also be used on the command line for generic data processing scenarios.

Benchmarking nsearch against other commonly used general purpose tools USEARCH (Edgar, 2010) and VSEARCH (Rognes et al., 2016) demontrates that nsearch can compete in performance with comparable state-of-the-art tools (both in run time and sensitivity). However, tools such as USEARCH and VSEARCH offer advanced features such as clustering which are not yet implemented in nsearch.

nsearch is available on GitHub under the permissive BSD-3-clause license at https://github.com/stevschmid/nsearch.

## Acknowledgements

This work was funded by the Bill and Melinda Gates Foundation grant OPP1151041 and the Swiss National Science Foundation (SNSF) through grant OP157065 to TRJ.

